# Age-Dependent Changes in the Progenitor Translatome Coordinated in part by *Tsc1* Increase Perception of Signaling Inputs to End Nephrogenesis

**DOI:** 10.1101/2020.03.12.989343

**Authors:** Eric Brunskill, Alison Jarmas, Praneet Chaturvedi, Raphael Kopan

**Affiliations:** Division of Developmental Biology, Department of Pediatrics, University of Cincinnati College of Medicine, Cincinnati, OH, USA

## Abstract

Mammalian nephron endowment is determined by the coordinated cessation of nephrogenesis in independent niches. Here we report that in young niches, cellular Wnt agonists are poorly translated, Fgf20 levels are high and R-spondin levels are low, resulting in a pro self-renewal environment. By contrast, older niches are low in Fgf20 and high in R-spondin, with increased cellular translation of Wnt agonists, including the signalosome-promoting Tmem59. This suggests a hypothesis that the tipping point for nephron progenitor exit from the niche is controlled by the gradual increase in stability and clustering of Wnt/Fzd complexes in individual cells, enhancing the response to ureteric bud-derived Wnt9b inputs and driving differentiation. We show *Tsc1* hemizygosity differentially promoted translation of Wnt antagonists over agonists, expanding a transitional (*Six2+, Cited1+, Wnt4+*) state and delaying the tipping point. As predicted by these findings, reducing Rspo3 dosage in nephron progenitors or Tmem59 globally increased nephron numbers *in vivo*.

## INTRODUCTION

Nephron number varies in humans 10-fold, ranging from 2×10^5^ to 2.5×10^6^ filtration units per kidney (Bertram et al., 2011; Hoy et al., 2005; Nyengaard and Bendtsen, 1992). Individuals with low nephron endowment are at high risk for developing hypertension, chronic kidney disease (CKD), and end stage renal disease (ESRD; (Barker et al., 2002; Barker, 1991, 1995; Barker and Bagby, 2005; Barker et al., 2006; Barker et al., 1990; Lackland et al., 2000; Rodriguez et al., 2005; Sayer et al., 1998; Sutherland et al., 2011). Infants born prior to 30 weeks gestational age undergo postnatal nephrogenesis for at most 40 days after birth (Rodriguez et al., 2005), and some/most nephrons formed after birth appear abnormal (Sutherland et al., 2011) and may not function properly. Currently, neonates born at 24 weeks have a survival rate of over 60% (Stoll et al., 2015), and are predicted to be at the extreme low end of nephron endowment and at high risk for early onset CKD and ESRD as compared to adults born at full-term (Crump et al., 2019; Haikerwal et al., 2020; Vikse et al., 2008; White et al., 2009). With the improvements in survival of extremely low birth weight and preterm infants, the negative implications of low nephron endowment and the associated economic and quality of life impacts will increase (Haikerwal et al., 2020). Understanding the mechanism(s) that control the synchronous cessation of nephrogenesis could be leveraged to develop interventions aimed at extending the process and increasing nephron endowment in at-risk populations, including premature and low-weight births.

Mammalian nephrogenesis depends on transiently amplifying nephron progenitors cells (NPCs) localized in a tight “cap” on the distal aspect of ureteric bud (UB) tips (Boyle et al., 2008; Kobayashi et al., 2008). The NPCs are sustained by juxtracrine Fgf20/9 signaling and reciprocal interactions between the NPCs and the UB, which secretes Wnt9b, a factor promoting both self-renewal and differentiation of NPCs (Costantini and Kopan, 2010; Dressler, 2009; Kopan et al., 2014). The balance between self-renewal and differentiation is regulated in part by stromal signals that may include Fat4 (Das et al., 2013) and/or Decorin (Fetting et al., 2014). Unlike the mesonephric kidney, which adds nephrons throughout adult life via long-lived stem cells (Diep et al., 2011), the metanephric kidney has a finite window during which nephrons are generated. In humans, NPCs are exhausted *en masse* by 34-36 weeks of gestation, with 60% of nephrons forming during the third trimester (Hinchliffe et al., 1991). In mice, nephrogenesis ends by P3 (Brunskill et al., 2011; Rumballe et al., 2011), and nephron endowment can range two-fold between different strains (Supplemental Figure S1A). Historically, the NPC population has been thought of as a transitory cell population that progresses from an uncommitted or naïve state (*Six2*^*+*^; *Cited1*^*+*^), to a committed, primed (*Six2*^*+*^; *Cited1*^*–*^; *Wnt4*^*+*^) state, to pre-tubular aggregates (PTA, *Six2*^*+*^; *Cited1*^*–*^; *Wnt4*^*+*^, *Pax8*^*+*^; *Fgf8*^*+*^) which will ultimately complete a mesenchymal-to-epithelial transition to form each nephron. However, this linear progression toward epithelium has been recently challenged with the observation that *Six2*^*+*^; *Cited1*^*–*^; *Wnt4*^*+*^ cells can revert back to the naïve, *Six2*^*+*^; *Cited1*^*+*^, *Wnt4*^*–*^ state (Lawlor et al., 2019).

Cessation of nephrogenesis is not due to an extrinsic trigger (e.g., loss of niche factors (Hartman et al., 2007)), to an intrinsic age-dependent change(s) affecting the progenitors’ ability to self-renew (e.g., (Kahn, 2011)), or to an altered cap mesenchyme (CM)/UB ratio (Short et al., 2014). Instead, heterochronic engraftment experiments demonstrated that when transplanted, older naïve mouse NPCs engraft in a young niche better than age-matched committed NPCs but more poorly than young, committed NPCs. Those that engraft and remain in the niche are surrounded by young, FGF20-expressing NPCs and can and contribute nephrons for up to twice the normal lifespan of a mouse NPC *in situ* (Chen et al., 2015). That led us to suggest a “tipping point” model controlling cessation: when most neighbors display a “young” signature (which includes Fgf20, a niche retention signal; (Barak et al., 2012; Chen et al., 2015)), NPCs gaining an “old” transcriptome signature persist in the niche. When the number of “old” cells among immediate neighbors rises above a hypothetical threshold, the model assumes all remaining progenitors will differentiate. *Tsc1* (hamartin) has emerged as one of the genes that might regulate the rate of progression to the tipping point, or the interpretation of the exit signal, since mice with NPCs hemizygous for this gene delayed cessation and have significantly more nephrons (Volovelsky et al., 2018). Despite these advances, the precise signaling mechanism(s) that drive the coordinated exit of NPCs from all nephrogenic niches at the time of cessation are not completely understood.

To identify molecular mechanism(s) controlling cessation of nephrogenesis, we leveraged two existing models with increased nephron endowment – *Six2-<u>G</u>FP<u>C</u>re<u>E</u>RT2*, (*Six2*^*GCE/+*^, herein *Six2*^*KI*^; (Combes et al., 2018; Kobayashi et al., 2008)) and *Six2*^*TGC*^;*Tsc1*^*+/Flox*^ (*Six2*^*TGC;Tsc1*^; (Volovelsky et al., 2018)) – and analyzed the transcriptome at two developmental timepoints (E14 and P0) of three genotypes: *Six2*^*KI*^, *Six2*^*TGC;Tsc1*^ and the control *Six2*^*TGC*^; (Kobayashi et al., 2008). We also analyzed the translatomes of control *Six2*^*TGC*^ and *Six2*^*TGC;Tsc1*^ NPCs. We discovered that the “old” (P0) cells enhanced the translation of components in all major signaling pathways, making these cells more acutely “aware” of their signaling environment. The analyses further established that the two advantageous mutations enhance nephrogenesis via temporally-disparate effects on the nephron progenitor population. *Six2* hemizygosity affects the young NPCs via a mechanism yet to be fully explored, whereas *Tsc1* hemizygosity differentially affected the relative abundance of key Wnt pathway components in older P0 NPC, lowering translation of agonists (*Rspo1, Rspo3*, and *Tmem59*) and elevating translation of antagonists (*Kremen1, Dkk1, Znrf3/Rnf43*) to reduce the relative strength of the Wnt9b signals received by these cells. *Tsc1* hemizygosity resulted in the accumulation of NPC co-expressing *Six2*^*+*^; *Cited1*^*+*^, *Wnt4*^*+*^, many of which appear to have been in the process of reverting to the naïve state. These observations led to the hypothesis that the tipping point is controlled by the gradual increase within individual cells in the stability and clustering of Wnt/Fzd complexes, increasing the magnitude of the response to Wnt9b inputs. We validated this hypothesis by demonstrating that *Rspo3* hemizygosity in NPCs (shortening the half-life of Wnt/Fzd complexes), or loss of the signalosome-promoter *Tmem59* (reducing avidity to Wnt9b and downstream signaling components; (Gerlach et al., 2018)) both increased nephron endowment. These findings provide a potential therapeutic framework for delaying cessation and increasing nephron endowment in at risk populations.

## RESULTS

### The cellular composition of the nephron progenitor niche is unaffected by mutations generating increased nephron numbers

Increased nephron numbers observed in separate genetic models could arise as a result of several different mechanisms. First, increased nephron endowment may reflect either an expanded or distinct, novel population of NPCs. Second, age-dependent change in the transcriptome (Chen et al., 2015) and/or metabolome (Liu et al., 2017) in these two models may unfold more slowly, enabling NPC to retain their “youthful” character relative to chronologically matched NPCs from the control strain, *Six2*^*TGC*^. Third, a molecular mechanism(s) unrelated to the relative age of the progenitors is involved. Predictions of all three hypotheses could be tested by analyzing the single-cell transcriptomes of NPC populations isolated from mouse kidneys of each of the three genotypes (Six2^TGC^, *Six2*^*TGC;Tsc1*^ and *Six2*^*KI*^) at two developmental time points (embryonic day 14 (E14) and postnatal day 0 (P0)). Single cell RNA-seq (scRNA-Seq) profiles generate sufficient markers to segregate even closely related cell types within tissues, enabling identification of NPC population(s) correlated with the increase in nephron numbers. As various different informatic packages exist, we follow best practices (Luecken and Theis, 2019) and when possible, use multiple pipelines/packages to seek bioinformatic consensus. To collect six scRNA-Seq datasets, one at E14 and one at P0 for each genotype, we dissociated kidneys for 5-7 min at 10°C with psychrophilic proteases (Adam et al., 2017) in an Eppendorf Thermomixer R, generated single cell suspensions, retained only samples with >85% viability (assessed by trypan blue staining), and submitted the pooled cells from several kidneys for sequencing on the 10X Genomics Chromium v2 platform. Our samples were enriched for cells in the renal cortical region due to the rapid, peripheral digestion of the tissue. The exclusion of a FACS-isolation step allowed us to 1) collect other relevant cell types in nephrogenesis and 2) reduce the potential introduction of cell stress signatures by extensive, high temperature sample processing or flow cytometry.

For the subsequent analyses, we used dimensionality-reducing software Seurat 2.4 and 3.0 (see extended experimental methods for detail). Unsupervised clustering using t-distributed stochastic neighbor embedding (t-SNE; (Macosko et al., 2015)) of the six samples defined between 14-18 transcriptionally distinct cell types across the genotypes and time points, representing various cell types derived from the developing kidney cortex including nephron progenitors (NPC), ureteric bud (UB), stroma (S), renal vesicle (RV), s-shape body (SSB), pretubular aggregate (PTA), proximal tubule (PT), and distal tubule cells (DT) (Figure 1A,B, Supplemental Figure S1B).

**Figure 1:**
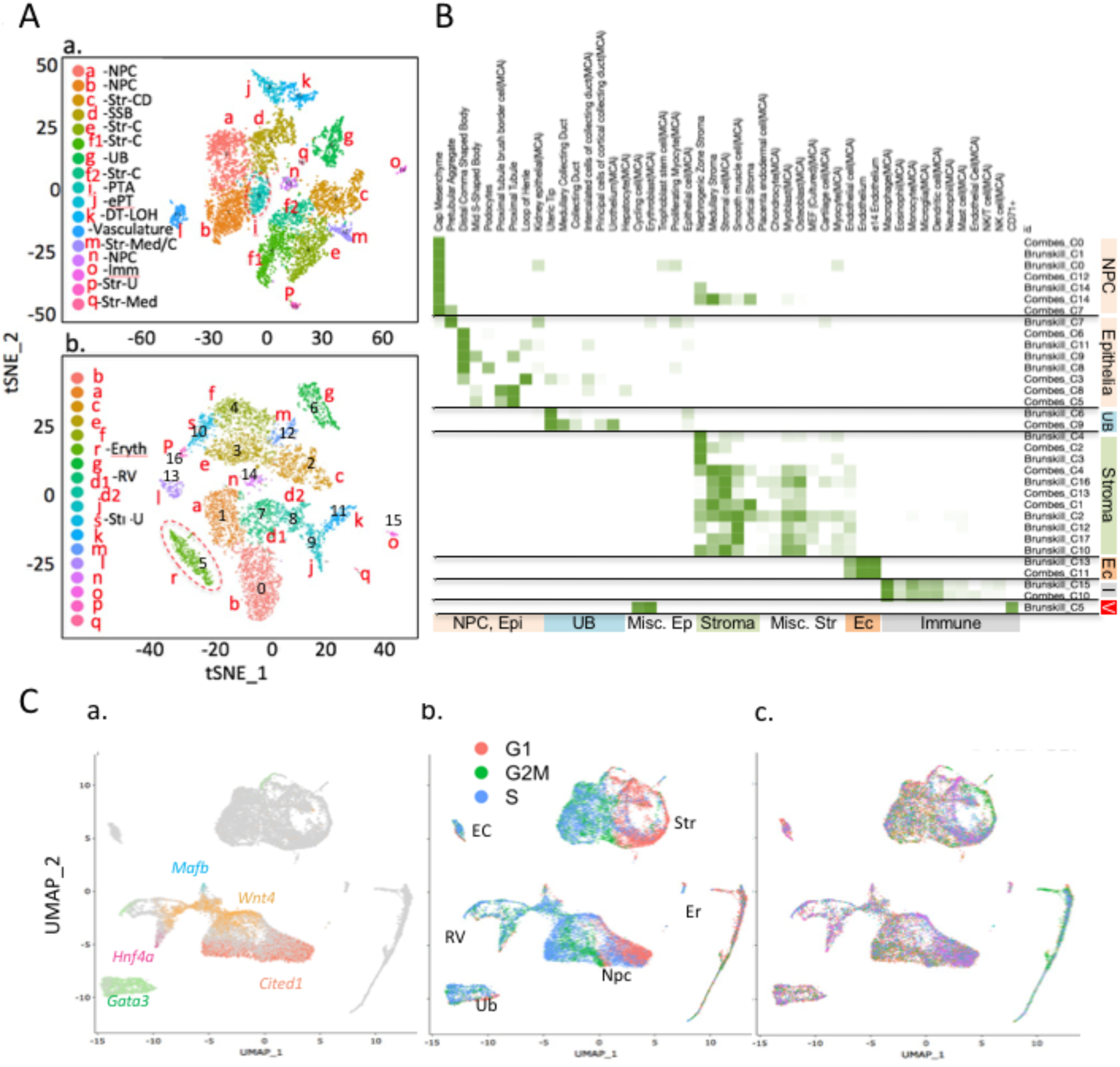
Cell populations identified in the developing kidney using different methodologies and different bioinformatic processes are indistinguishable. (Aa) Six2^KI^ data, transcripts included after cell cycle removal; (Ab) same as (Aa), filtered as in Combes et al, 2019. Annotation of clusters based on similarity to Combes et al. Note cluster d in (Aa) was split in (Ab), and cluster f in (Ab) was split in (Aa). (B) GO Elite analysis of Combes’ and our Six2^KI^ against global datasets. Note cortical bias in our clusters. (C) UMAP clustering of all six samples. Gene expression patterns were merged in photoshop (Ca) to illustrate the temporal progression from naïve (Cited1+, right) to epithelial (Gata3+, Hnf4a+, Mafb+) at left. Wnt4 marks transitional “primed” NPC. Cell cycle states and cluster identities are shown in (Cb), and the genotype contribution to each cluster in (Cc).

To confirm that we captured all potential cortical cell types found in the developing kidney, and to evaluate how different bioinformatic parameters could potentially mask or reveal the presence of novel cell clusters, we cross-referenced our data with other mouse kidney scRNA datasets. To accomplish this, we performed a similarity analysis by comparing our E14 *Six2*^*KI*^ data to an existing *Six2*^*KI*^ dataset (Combes et al., 2019) using the Gene Set Enrichment Analyses (GSEA) algorithm. Combes et. al. analyzed wildtype and Six2 hemizygotes processed via dissociation at 37°C, performed multiple independent scRNAseq runs to control for batch effects, and filtered out ribosomal, mitochondrial, and lncRNA genes in their analytical pipeline (using Seurat 2.4). We performed a pairwise analysis between the two datasets using bioinformatic parameters which either included ribosomal, mitochondrial and lncRNAs or filtered-out these gene components. This analysis demonstrated that all three *Six2*^*KI*^ datasets (Combes’, and ours analyzed with two different filter sets) were highly correlated [Figure 1A, B, Supplemental Table S1 (our unfiltered gene set vs. Combes’), Supplemental Table S2 (our filtered gene set vs Combes’)]. We developed a script to extract the FDR score assigned by the GSEA algorithm for each pairwise comparison between our clusters and those described by Combes et al. in their Table S1 (Combes et al., 2019) and plotted the scores (Supplemental Figure S1C). Importantly, while the methodological differences changed the relative rank of individual marker genes, splitting one cluster (cortical Stroma, Str-C, Figure 1Ab, *f* into Figure 1Aa, *f1*&*f2*) and fusing another (S-shaped body, SSB, Figure 1Aa, *d* into Figure 1Ab, *d1*&*d2*)), all (and only) the cell identities assigned by Combes et al. were present in each of our samples (Supplemental Figure S1C). These analyses confirmed that our scRNA methodology captured a comparable collection of developing kidney cells. As an additional, unbiased test of similarity we used GO-Elite (www.genmapp.org) to identify previously annotated cell types most highly correlated with similarly filtered clusters from Combes and our data (Figure 1B). This analysis correctly identified the cortical bias in our scRNA dataset (e.g., lack of mature proximal tubule cells, ductal stalk cell types) resulting from our sample preparation method, and showed that the NPC-Str cluster (Combes C14) was a minimal contributor in our data, consistent with it being a technical artifact as proposed by the authors. Taken together, these analyses demonstrate that despite technical differences in sample preparation and bioinformatic biases introduced through alternative filtration methods, the datasets are highly correlated and cortically-derived cell types found in developing kidneys are well represented in our data.

We proceeded to investigate the possibility that a unique cell population distinguished the high nephron number lines (*Six2*^*K*I^ and *Six2*^*TGC;Tsc1*^) from the control (*Six2*^*TGC*^). We pooled all six samples and performed an integration analysis using the Seurat 3.0 UMAP algorithm (Stuart et al., 2019) to visualize clusters which emphasized global similarities. This analysis identified several super clusters, characterized as an NPC cluster, stromal cluster, ureteric bud (UB) cluster, and minor clusters for erythrocytes, immune cells and endothelial cells (Figure 1C). Notably, UMAP correctly clustered the NPC along a proximo-distal developmental axis, with the naïve *Six2+, Cited1+* cells at one pole of the cluster, followed by a transitional population (*Six2+, Wnt4+*), and segregating into epithelial precursors of proximal tubule (PT, *Hnf4a+*), distal tubule (DT, *Gata3+*) and podocyte (Pod, *Mafb+*) at the opposite pole (Figure 1Ca, Figure 2A-D). Most of the naïve NPCs were in G1 (Figure 1Cb, 2D). Importantly, assigning a unique color to each genotype and developmental time point (palette in Figure 2L) reveals that unlike erythrocytes, the majority of which were contributed by the E14 *Six2*^*TGC;Tsc1*^ sample (green), no sample dominated a particular area of the NPC super cluster (Figure 1C, 2A). These analyses allowed us to conclude that the differences between the two strains generating more nephrons did not reflect overabundance of any particular NPC subtype. Analysis of stromal and UB clusters will be reported elsewhere.

**Figure 2:**
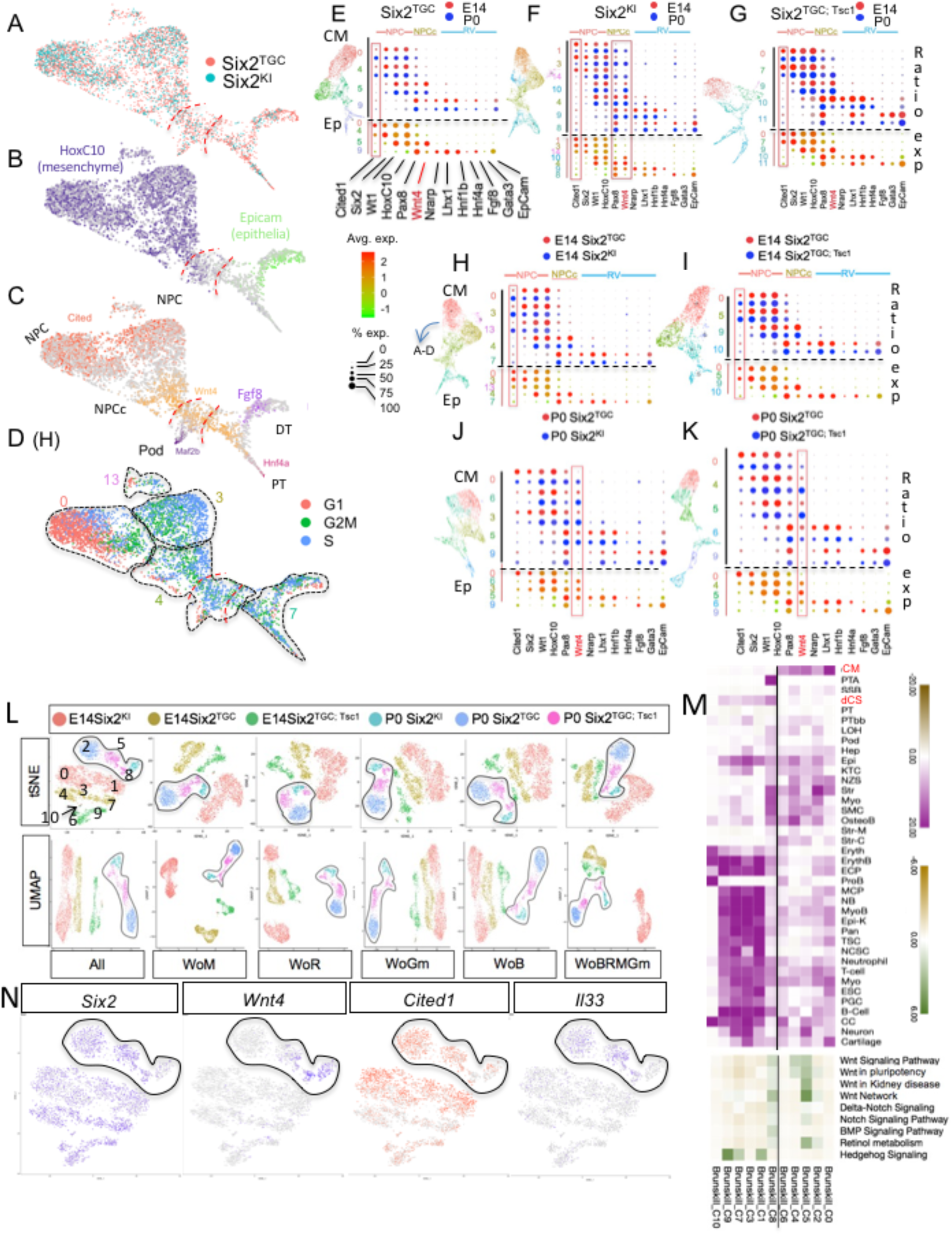
Pairwise UMAP integration analyses indicate that Six2KI and Six2^TGC;Tsc1^ P0 NPC are less “youthful” in character than control Six2^TGC^. (A-D) NPC cluster clipped from UMAP integration analysis of P0 Six2^KI^ and Six2^TGC^ (shown in H). (A) Distribution of cells across the NPC cluster, assigning a unique color to each genotype and developmental time point (B) Expression of *Hoxc10* (mesenchymal) and *Epcam* (epithelia) merged in photoshop illustrates MET across the cluster. (C) Gene expression patterns were merged in photoshop to illustrate the temporal progression from naïve (*Cited1+*, left) to epithelial *(Gata3+, Hnf4a+, Mafb+*) to the right. (D) Cell cycle status of each NPC sub-cluster (traced and numbered). Note that these numbers are represented on the Y axis of panels E to K. (E-G) Pair-wise integration analysis of E14 and P0 NPC for each genotype. On the left, the cluster is shown with colors corresponding to cluster numbers on the Y axis, arranged from Naïve (top) to epithelial (bottom). On the X axis, selected genes are arranged from left (naïve) to right (epithelial). For each genotype, the fraction of cells expressing a given gene is reflected in the diameter of red (E14) or blue (P0) circles. The level of expression is plotted below in red (high) and green (low) circles. Red box highlights Cited1 expression. (H-K) Pair-wise integration analysis of E14 (H-I) or P0 (J-K) NPC for control Six2TGC (red circles) vs. Six2^KI^ (blue circles, H; J) or vs. Six2^TGC^; Tsc1 (blue circles, I; K). Red box emphasizes Wnt4 expression. (L) Clustering all NPCs across six samples using tSNE and UMAP algorithms. All denotes no filter; WoM- without mitochondrial genes; WoR- without ribosomal genes; WoGm- without gene model transcripts; WoB- without blood transcripts; WoBRMGm- without all these entities. P0 clusters are traced. See text for detail. (M) Go ELITE analysis of NPC clusters for cell type (purple) or selected pathways (Green). (N) Selected gene expression patterns across “All” tSNE clusters.

To increase the resolution and find possible gene expression signatures that might be masked by analyzing all six samples at once, we performed pair-wise integration analysis. First, we compared E14 to P0 cells for each genotype. For each NPC super cluster, we analyzed the fraction of cells expressing a given gene (reflected in the diameter of red (E14) or blue (P0) circles), and the level of expression (red/green circles). We selected genes with graded expression representing naïve NPC (*Cited1, Six2, Hoxc10, Wt1*), committed or transitional NPC (*Six2, Wt1, Wnt4, Pax8*), PTA (*Nrarp, Lhx1, Hnf1b*) and the emerging epithelial linages (*Epcam, Hnf4a, Fgf8, Gata3*). Intriguingly, we noted that in the control *Six2*^*TGC*^ strain, more older cells expressed *Cited1* than young cells (Figure 2E). Inversely, more E14 *Six2*^*KI*^ cells expressed *Cited1* than P0 *Six2*^*KI*^ (Figure 2F), whereas E14 and P0 *Six2*^*TGC;Tsc1*^ NPC contributed equally to *Cited1+* population (Figure 2G). Next, we performed pairwise comparisons of each high nephron-producing NPCs (blue circles, (Figure 2H-K) to its age-matched *Six2*^*TGC*^ control (red circle, Figure 2H-K). Surprisingly, E14 *Six2*^*KI*^ and *Six2*^*TGC;Tsc1*^ resembled P0 *Six2*^*TGC*^ cells as they contributed more *Cited1*-expressing NPC (Compare Figure 2H, I with 2E). At P0, Six*2*^*KI*^ and *Six2*^*TGC;Tsc1*^ contributed most of the *Wnt4*-expressing cells (Figure 2J,K). This is strikingly inconsistent with the hypothesis that NPCs from *Six2*^*KI*^ and *Six2*^*TGC;Tsc1*^ retain a youthful, naïve character relative to control.

### *Six2, Tsc1* hemizygous NPCs produce more nephrons yet appear older and more differentiated than controls at P0

In our previous report, we used the Fluidigm C1 system to establish that transcriptional signatures distinguish “old” (E18.5, P0) from “young” (E12, E14.5) NPC, and the most notable changes were increased mitochondrial and ribosomal signatures in E18.5 and P0 NPC compared to E12.5-E14 NPCs (Chen et al., 2015). To address the relative age of *Six2*^*KI*^ and *Six2*^*TGC;Tsc1*^ NPC, we pooled all the NPC clusters identified based on their GSEA FDR scores (Supplemental Figure S1C, red boxes). Each cell was provided with a unique identifier and the pooled NPC population re-clustered using Seurat 3.0 under six filtering permutations (Figure 2L): no filter (all transcripts included), without mitochondrial genes (WoM), ribosomal genes (WoR), gene model transcripts (WoGm), blood transcripts (WoB) or excluding all of the above commonly filtered gene sets (WoBRMGm). Regardless of the filter or clustering algorithm (t-SNE or UMAP) applied, six superclusters, each corresponding to a unique genotype/age combination, emerged. All P0 cells clustered together and separately from E14 cells. *Six2*^*KI*^ E14 segregated distinctly away from the other E14 samples, most notably when WoR, WoM and WoBRMGm filters were applied. This analysis confirmed a key finding underlying the “tipping point” hypothesis (Chen et al., 2015): that NPC transcriptomes are indeed altered over the ten days of murine metanephrogenesis, including but not limited to the respiratory and translational machinery components. Furthermore, this analysis showed that the effects of *Six2* and *Tsc1* hemizygosity are profound enough to distinguish these cells not only from controls at both timepoints, but also from each other. This suggests that multiple pathways and mechanisms may be responsible for the gene expression differences that separate these genotypes.

To further identify global trends from our datasets, the top markers of each cluster were analyzed using the AltAnalyze GO-Elite function, which examines Gene Ontology overrepresentation covering gene, disease and phenotype ontologies, multiple pathway databases, biomarkers, transcription factor and microRNA targets (Zambon et al., 2012). This analysis assigned cell types to each NPC cluster that was most highly correlated with its gene signature (Figure 2M, purple, and Supplemental Table S4). Interestingly, this analysis resolved the NPC clusters into two distinct groups: A Cap Mesenchyme group (the E14 clusters C0, C2, C4, and the P0 clusters C5 and C6), and an early epithelia group. All the P0 *Six2*^*KI*^ NPC, together with a few *Six2*^*TGC;Tsc1*^ cells, resembled pretubular aggregate cells (PTA, cluster C8; cluster numbers shown in Figure 2L). The rest of the NPC clusters (C1, C3, C7, C9 and C10) were classified as early epithelia, resembling distal comma shape body (Figure 2M, dCS), and contained transcripts enriched for an eclectic collection of stem cells, epithelial cell types, and immune cells. Importantly, GO-Elite wiki-pathway analysis identified the highest enrichment for the WNT signaling (*Fhl2, Fzd2, Map1b, Tcf4, Wnt4*) and Retinol metabolism pathway (*Aldh1a, Crabp1, Rbp1*) in *Six2*^*TGC;Tsc1*^ cells (Cluster C5) compared to the other NPCs (Supplemental Figure S2). The entire cap mesenchyme (CM) group was also enriched for ribosomal proteins, BMP and Notch signaling, whereas the non-CM NPCs were enriched for ontologies related to Hedgehog signaling, DNA replication, and one carbon metabolism (Figure 2M, Supplemental Figure S2). Electron transport and oxidative phosphorylation pathways where enriched in non-CM E14 *Six2*^*TGC*^ clusters and P0 *Six2*^*TGC;Tsc1*^ CM, inconsistent with reduced respiration driving increased nephron endowment in the latter population (Liu et al., 2017). Given the high levels of *Cited1* at E14 in *Six2*^*KI*^ (Figure 2F), distinct clustering of E14 *Six2*^*KI*^ cells away from age-matched cells, we surmise that the increased nephron endowment in *Six2*^*KI*^ might be driven by a mechanism acting earlier in nephrogenesis, before P0. By contrast, our analysis suggests that this mechanism was acting late in *Six2*^*TGC;Tsc1*^. Notably, this analysis did not uncover evidence to support the hypothesis that P0 *Six2*^*TGC;Tsc1*^ progenitors were more “youthful” than controls. Quite to the contrary, like the *Six2*^*KI*^ NPC, P0 *Six2*^*TGC;Tsc1*^ cells contained higher levels of Wnt4 mRNA and evidence of Wnt pathway activity (Figure 2M), associated with mesenchymal to epithelial transition (MET) and differentiation. Unlike P0 *Six2*^*KI*^ NPC, they were still clearly CM cells.

### Tsc1 hemizygous nephron progenitors contain a significant population of *Six2+, Cited1+, Wnt4+* cells

Our analysis suggested that the Wnt signaling pathway may be altered in our *Tsc1* hemizygote model. A closer examination of Wnt signaling gene expression in cap mesenchyme clusters suggested that *Wnt4* levels were graded in P0 clusters: low in Six2TGC (C2), more in Six2TGC^;*Tsc1*^ (C5) and high in *Six2*^*KI*^ (C8, Figure 2N). *Cited1* and *Il33* expression were graded in the opposite direction, highest in P0 *Six2*^*TGC*^ (scale favors display of the cells with expression above median). To more carefully analyze the behavior of NPCs of different ages, we re-clustered NPCs with RNA Velocity and scVelo algorithms, which analyzes the ratio of spliced to un-spliced transcripts to predict the trajectory of cells within the UMAP multidimensional landscape (see methods for detail). Both algorithms segregated naïve cap mesenchyme cells (*Six2+, Cited1+, Wnt4–*) into three main lobes, away from the committed NPC (*Six2+, Cited1–, Wnt4+*; Figure 3A). Cells selecting distal tubule fate (*Gata3+*) segregated to the opposite pole of cells selecting the proximal tubule fate (*Hnf4a+*), while podocyte precursors (*Mafb+*) remained embedded within committed NPC (Figure 3B). Layering the genotype information of each cell revealed that the lobular structure separated the E14 *Six2*^*KI*^ into one lobe, the E14 *Six2*^*TGC*^ and *Six2*^*TGC;Tsc1*^ into an intermediate lobe, and the P0 *Six2*^*TGC*^ and *Six2*^*TGC;Tsc1*^ into the third lobe (Figure 3B). Interestingly, and consistent with our interpretation, P0 *Six2*^*KI*^ cells remain embedded with the committed NPC. Notably, very little overlap was detected between *Cited1* and *Wnt4* expression in all E14 clusters, but considerable overlap was evident in the P0 *Six2*^*TGC*^ and *Six2*^*TGC;Tsc1*^ clusters (see tracing of expression in Figure 3A,B). To quantify this, we calculated the fraction of cells with the signature *Six2+, Cited1+, Wnt4+* in each genotype by age and noticed that whereas the control E14 *Six2*^*TGC*^ cluster contained less than 1% of cells with this signature, all P0 clusters contained ∼4% or more cells. Strikingly, 12.5% of cells in the P0 *Six2*^*TGC;Tsc1*^ cluster display this combined *Six2+, Cited1+, Wnt4+* profile. This observation calls attention to the possibility that more *Six2*^*TGC;Tsc1*^ cells are migrating back to the naïve territory, as reported recently for the Wnt4+ population in developing kidneys (Lawlor et al., 2019). Velocity embedding revealed that indeed, *Wnt4+ Six2*^*TGC;Tsc1*^ cells tended to contain un-spliced (recently transcribed) *Cited1* RNA (Figure 3D, E). These analyses shifted our focus to the Wnt pathway, with the caveat that cellular RNA in general, and scRNA in particular, are poor predictors of the proteome and thus are rather limited as a foundation for developing and investigating mechanistic hypotheses.

**Figure 3:**
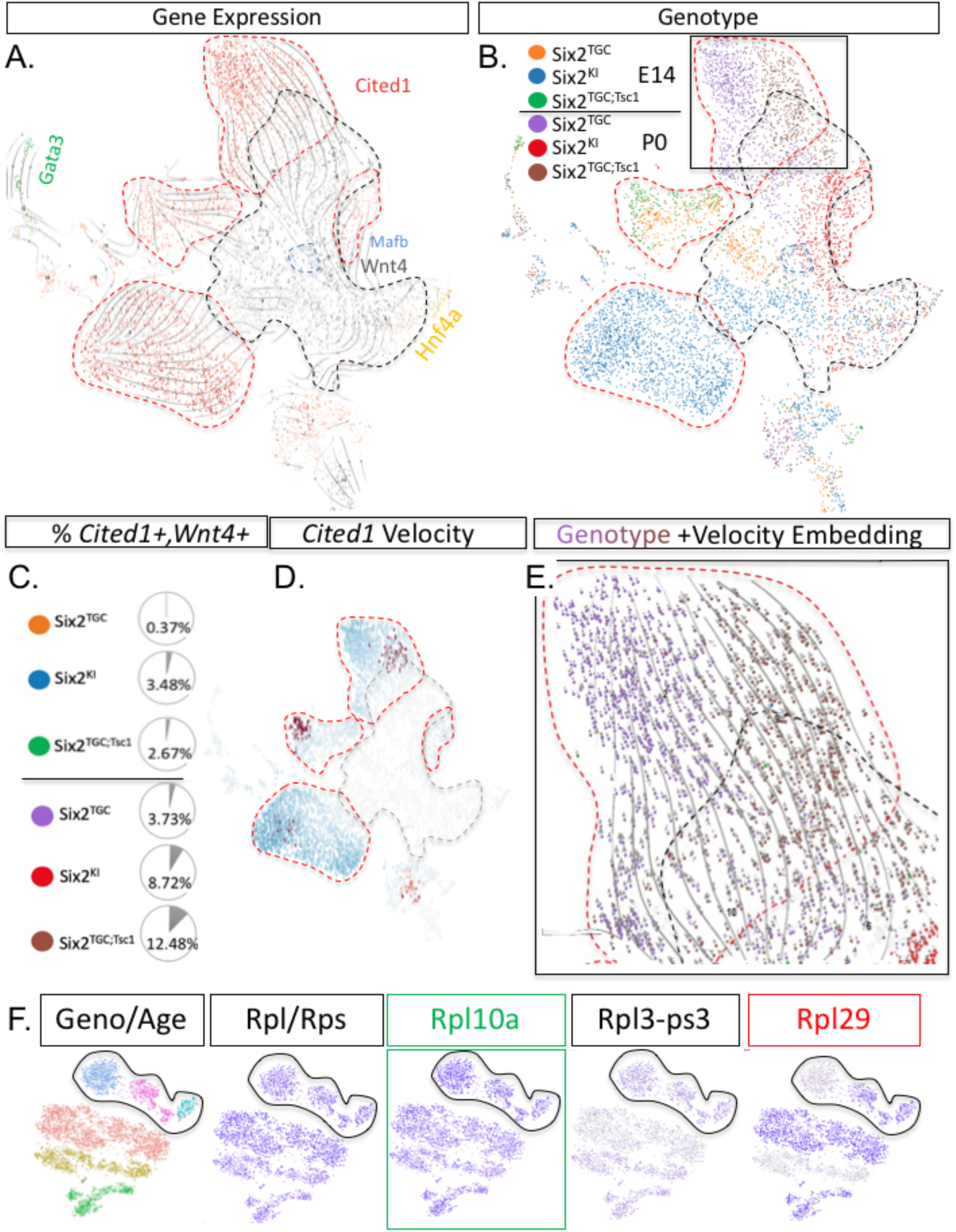
RNA velocity analysis identifies an increase in a population with the combined *Six2*^*+*^, *Cited1*^*+*^, *Wnt4*^*+*^ expression profile. (A). RNA Velocity-clustered NPCs visualized with scVelo. Shown are gene expression domains for *Cited1, Wnt4*, and markers for PT (*Hnf4a*), DT (*Gata3*) and POD (*Mafb*). The faint black lines are velocity arrows. (B) The same plot as in (A), overlaid with color-coded genotype/age information. *Cited1* expression traced with red dashed line, *Wnt4* expression traced in dashed back line. (C) The fraction of *Six2+* cells with the signature *Cited1+, Wnt4+* in each genotype by age. (D) The same plot as in A, colorized for cells with spliced transcripts (blue) and those accumulating newly transcribed *Cited1* transcripts (brown). (E) Close up of the P0 Six2^TGC^ (purple) and P0 Six2^TGC, Tsc1^ (brown) cells with the embedded velocity (gray arrowhead). Arrow size reflects amount of *Cited1* transcripts and the direction the cell differentiates. Note *Wnt4*^*+*^ cells transitioning to *Cited1* cells, not vice versa. (F) Expression of selected ribosomal transcripts in the tSNE unfiltered clusters as presented in Figure 2L.

### Translatome analysis uncovers a translation bias in *Tsc1* hemizygous NPCs which could impact the interpretation of Wnt9b signals

Given the limitations of scRNA-seq (e.g. discordance with the protein state of the cell (Kristensen et al., 2013), and dropout/undetected transcripts), we shifted to an alternative strategy of Translating Ribosome Affinity Purification (TRAP; (Doyle et al., 2008)) which has proven to provide a translation profile with high concordance to the proteome. We examined our scRNA-Seq data to ensure that ribosomal protein Rpl10a was indeed expressed in all cells at both time points, as we noted that there was some ribosomal composition variability among the strains (see Figure 3F, Rpl29, (Shi et al., 2017)). Mice engineered to contain the ribosomal protein EGFP fusion (EGFP/Rpl10a) with a floxed stop cassette (JAX Stock #024750) in front of the powerful CAGGS promoter (Miyazaki et al., 1989) were crossed to *Six2*^*TGC*^ or *Six2*^*TGC;Tsc1*^ to activate expression of EGFP/Rpl10a in the NPC and their descendants. To analyze the translatome, cells were sorted for expression of GFP^HI^ (Six2^TGC^ and EGFP/Rpl10a, the latter also expressed in nephron epithelia) and Itg8a^HI^ (NPC, a few stromal cells), with negative selection for Pdgfra (stromal marker). The ribosomes were pulled down from E14 control (*Six2*^*TGC*^) and P0 animals of both genotypes (*Six2*^*TGC*^ or *Six2*^*TGC;Tsc1*^) and the RNA was isolated as described (Doyle et al., 2008). Bulk sequencing revealed several important insights regarding the changes occurring in the NPC during nephrogenesis.

Analysis of the RNA-Seq revealed significant differences between the E14 and P0 translatomes. For example, GO term analysis of differentially expressed genes identified a strong signature of cell cycle regulators in E14 *Six2*^*TGC*^, and translation of signaling pathways components (GO:0009719, GO:0071495, GO:0007167, p<10^−14^, FDR<10^−10^) that were significantly enriched in P0 *Six2*^*TGC*^ (Figure 4A, Supplemental Table S5), a trend becoming more pronounced in P0 *Six2*^*TGC;Tsc1*^. Most GO terms specifically enriched in P0 *Six2*^*TGC;Tsc1*^ were related to neurogenesis, due to the prominence of genes associated with axon pathfinding. Interestingly, one GO term described a cohort of 74 motility-related transcripts (GO:004887 p<10^−8^, FDR <10^− 5^) that were differentially translated in P0 *Six2*^*TGC;Tsc1*^ relative to control P0 NPCs (Supplemental Table S5).

**Figure 4:**
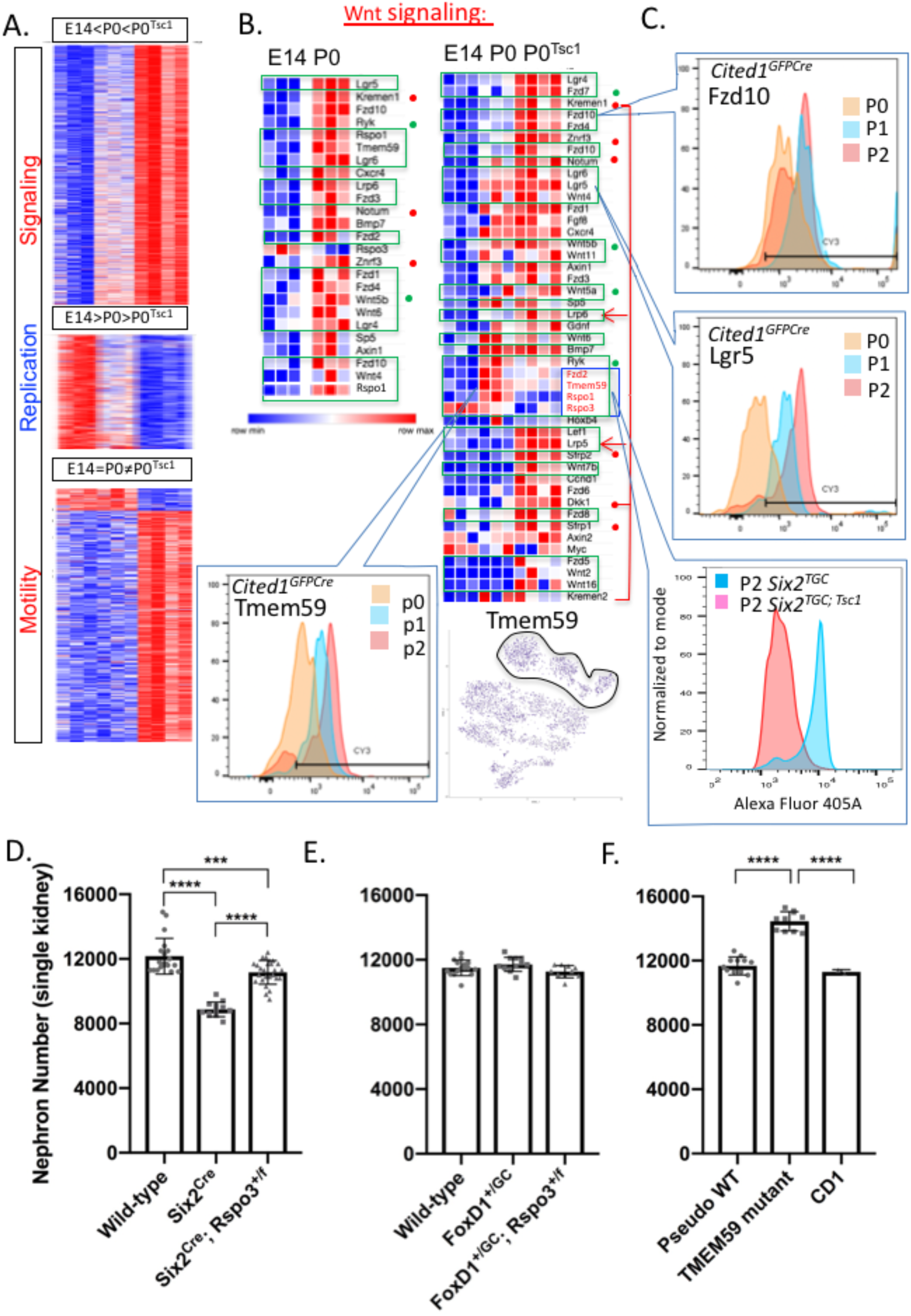
Translatome analysis and genetic validation models identify Wnt signaling as a mediator of the *Tsc1* enhanced nephrogenesis phenotype. (A) Differentially translated transcripts in sorted NPC isolated from E14, P0 Six2^TGC^ and P0 Six2^TGC; Tsc1^. The data are shown as relative change by row, with full expression data shown in Table S5. Top representative GO term for each subset of genes is shown to the left (see Supplemental Fig 2 for breakdown of signaling pathways). (B) On the left, a comparison of differentially translated Wnt pathway transcripts between E14 and P0 Six2^TGC^ highlights gains in signal reception machinery (agonists in green boxes, antagonists marked with red dots, non-canonical signaling marked with green dots). On the right, comparing all three samples, note increase in antagonists (beyond the change observed between E14 and P0 control cells) and decrease in agonists in P0 Six2^TGC; Tsc1^ (C) Specific proteins were selected for validation by flow cytometry, with good concordance with the translatome data. Note the TMEM59 levels increase with developmental age (left) and decrease in *Tsc1* hemizygotes (right). Note that unlike *Cited1, Wnt4* or *Il33, Tmem59* transcripts do not change with age (middle). (D-F) To validate the prediction that nephron number is dependent on a Wnt9b signal threshold, which progressively rises as translation of key components is increasing, we analyzed nephron numbers in adult (≥<u>P28)</u> mice with (D) NPC hemizygous for *Rspo3*. (E) stroma hemizygous to *Rspo3* and (F) global deficiency for TMEM59. Two-tailed unpaired t tests were performed in GraphPad Prism version 8 to evaluate statistical significance of single kidney nephron numbers. Both kidneys were counted for each animal with males and females represented. *** P ≤ 0.001; **** P ≤ 0.0001.

Given the evidence outlined thus far, we focused our attention on signaling pathways. Consistent with the tipping-point hypothesis, translation of components in all signaling pathways were elevated in older P0 *Six2*^*TGC*^ NPCs (Supplemental Figure S3A). Notably, most, but not all transcripts were translated more robustly in P0 *Six2*^*TGC;Tsc1*^ relative to P0 *Six2*^*TGC*^ NPCs, again consistent with finding that these progenitors are closer to an epithelial transition (elevated Hippo and Notch signaling) and less “youthful” overall. Among the transcripts that were translated more efficiently in *Six2*^*TGC*^ at P0 were many Wnt agonists (green boxes, green dot marks non-canonical agonist). By contrast, in P0 *Six2*^*TGC;Tsc1*^ transcripts of Wnt antagonists (red dots) were significantly more abundant in the translatome. They included *Dkk1, Kremen1*, and *Kremen2*, which together target the Lrp co-receptors for degradation (arrows in Figure 4B), and *Znrf3/Rnf43* which, in the absence of R-spondin/Lgr5 proteins targets the Wnt/Fzd complex for degradation (all marked with red dots in Figure 4B, (Nusse and Clevers, 2017)). Importantly, key Wnt agonists were among the transcripts translated less efficiently in P0 *Six2*^*TGC;Tsc1*^ relative to P0 *Six2*^*TGC*^ NPCs (blue box, Figure 4B): *Rspo1*, and *Rspo3*; required to engage *Znrf3/Rnf43* and prolong the half-life of the Wnt/Fzd complex, and *Tmem59*, coding for a molecule thought to cluster Fzd proteins thus increasing avidity/signal strength of WNT signaling (Gerlach et al., 2018). This combination of increased antagonists and decreased agonists was unique to the Wnt signal transduction pathway (Supplemental Figure S3), suggesting the compelling hypothesis that hamartin/Tsc1 regulates the strength of the Wnt9b signal by affecting the translation of key transcripts, and the impact on nephrogenesis in *Tsc1* hemizygotes might be due to reduced Wnt signal input below the threshold required for completion of MET, increasing the fraction of *Wnt4+, Cited1+* NPC, and delaying NPC exit from the niche and cessation of nephrogenesis.

To test this hypothesis, we first asked if increased transcript abundances in the translatome corresponded to increased abundance of Lgr5, Fzd10, and Tmem59 proteins. Flow cytometry of dissociated, fixed and permeabilized NPCs confirmed the concordance of the intracellular pool with the translatome for all three, including a substantial decrease in intracellular Tmem59 abundance in P2 *Six2*^*TGC;Tsc1*^ relative to P2 *Six2*^*TGC*^ (Figure 4C). Encouraged by these results, we pursued functional validation of our findings in genetic models. Nephron number was determined by the acid maceration method (Peterson et al., 2019) in young adult (≥P28) mice. We analyzed nephron numbers in *Six2*^*TGC*^; *Rspo3*^*+/f*^, to ask if a partial reduction in the levels of R-spondin 3, with an anticipated decrease in Wnt/Fzd complex stability, would impact nephron numbers. As an additional control we examined *FoxD1*^*Cre*^; *Rspo3*^*+/f*^, to ask whether *Rspo3* produced in the stroma affected nephron numbers to the same degree as that produced by the NPC. Notably, the *Six2*^*TGC*^ allele generates an approximate 30% deficit in nephron number compared to wild type, while *FoxD1*^*Cre*^ does not impact nephron endowment. As shown in Figure 4D, removal of one *Rspo3* allele in NPC resulted in a significant increase in nephron number, similar in magnitude to that seen in *Six2*^*TGC;Tsc1*^ mice. In contrast, no change in nephron numbers was detected when the *Rspo3* allele was removed from the stroma, indicating that this factor, like Fgf20, acts in a juxtracrine manner (Figure 4E). To address the role of Wnt signalosome formation, we designed sgRNAs to introduce a mutation within Exon4 of the *Tmem59* gene. Fertilized CD1 eggs were injected with dCas9 and gRNA, and 18 offspring were genotyped by cloning PCR products and sequencing 18 clones to verify the composition of both *Tmem59* alleles (Table S6). From this initial cohort, two homozygous mice, two pseudo-wildtype (harboring a six base pair in-frame deletion) and one heterozygote wildtype adult were analyzed. Subsequently, three additional *Tmem59* mutant offspring were analyzed. Wild-type CD1 nephron number data is also presented as a control for the CRISPR targeting process. As seen in Figure 4F, loss of *Tmem59* resulted in a significant increase in nephron numbers, confirming that reducing Wnt signal input in NPC enhances nephrogenesis.

## DISCUSSION

Transcriptome analysis at the single cell level is a powerful new tool applied effectively to build detailed maps of cellular constituents and their developmental trajectories in various tissues. While multiple studies have generated single-cell transcriptomic data for mouse (Adam et al., 2017; Brunskill et al., 2014; Combes et al., 2019; Magella et al., 2018) and human (Lindstrom et al., 2018; Wang et al., 2018) kidney development, the overwhelming focus in these early phases has been on enumerating the cell types compromising the kidney, with mechanistic exploration delegated for future investigation.

In combination with TRAP analysis, scRNAseq analysis of samples from two developmental timepoints across three genotypes, and the metanalysis of our data in the context of prior datasets, we reached several important conclusions. First, we observed that nephron endowment can be influenced by either molecular changes occurring early in nephrogenesis (as in *Six2* hemizygotes) or late in nephrogenesis (as in *Tsc1* hemizygotes), likely operating via distinct mechanisms. Second, a plausible mechanism impacting the timing of cessation is age-dependent increased translation of receptors, coreceptors, ligands, and other signal-reception enhancing proteins across all the ten pathways we examined (Notch, Wnt, FGF, BMP, TGF, HH, YAP, RA, PDGFR, VEGF). Thus, there is no need to change the niche signaling milieu – a key component in the tipping point appears to be increased “awareness” in older NPCs to their existing signaling environment. These translational changes may work synergistically or independently from the Lin28/Let7 axis (Yermalovich et al., 2019) or metabolism (Liu et al., 2017). Third, whereas “young” NPC in the Six2^TGC^ strain are indeed either naïve (*Six2*^*+*^, *Cited1*^*+*^, *Wnt4*^*–*^) or “primed” (*Six2*^*+*^, *Cited1*^*–*^, *Wnt4*^*+*^) (Mugford et al., 2009), ∼3% older NPC in the same strain are reverting from a “primed” to naïve state (Lawlor et al., 2019), identified by a triple positive (*Six2*^*+*^, *Cited1*^*+*^, *Wnt4*^*+*^) state due to the presence of unspliced, new *Cited1* transcripts. Fourth, reduction in *Tsc1* dose resulted in enhanced translation across all signaling pathways, and NPC appeared “older” and more PTA-like than control. This effect may well be dependent on enhanced mTor activity, although this has not been tested. Fifth, unexpectedly, *Tsc1* hemizygosity also resulted in asymmetric translation of several transcripts, including Wnt singling components, with key agonists (R-spondin proteins, Tmem59) translated less efficiently and key antagonists (Dkk1/Kremen, Znrf3/Rnf43) translated more efficiently. How hamartin impacts this asymmetry is a topic of active investigation.

It has been appreciated for over a decade that a UB-derived Wnt9b canonical signal is required for both maintenance of self-renewal capacity of NPC as well as for their differentiation (Carroll et al., 2005; Karner et al., 2011). Recently, work in cells derived from E10.5 mesonephric mesenchyme identified a role for Rspo1, Lrp6 and Fzd5 in enabling Wnt9b responsiveness as cells transitioned from the intermediate mesoderm to the metanephric mesenchyme identity (Dickinson et al., 2019). It was further demonstrated that low signaling input promotes self-renewal and high signaling input promotes differentiation (Ramalingam et al., 2018). Our findings explain how Wnt9b signal strength varies to control both decisions, changing over time: a cell intrinsic and progressive increase in translation leads to increased stability (more Rspo, Lrp, Lgr proteins) and clustering (via Tmem59) of Wnt/FZD complexes. This enables a cell to receive a more sustained and/or stronger Wnt9b signal. As more cells to switch to stable, clustered Wnt/Fzd complexes, coupled with additional changes yet to be characterized, the entire population switches from self-renewal to differentiation. We validated this by demonstrating that reduced Tmem59 indeed elevated nephron numbers. However, how can we reconcile this cell intrinsic process with the observation that “old” cells in a “young” neighborhood continue to self-renew (Chen et al., 2015)?

Since modulation of *Rspo3* dosage within the FoxD1 linage did not impact nephron number but *Rspo3* hemizygously in NPC did, we speculate that one mechanism explaining how “young” cells enable “old” cells to remain in the niche (Chen et al., 2015) is by providing Fgf20 and diluting Rspo1/3 below the threshold required to send NPC to the exit. Note that Fgf9 production is elevated at P0, perhaps because Fgf20 alone cannot counter the increase in Wnt9b signal received. The 3-fold increase in *Six2*^*+*^, *Cited1*^*+*^, *Wnt4*^*+*^ cells in kidneys with *Tsc1* hemizygous NPC also reflects an environment in which Rspo1/3 are reduced, and a cellular context with reduced clustering. Together, the data suggests that a very precise threshold needs to be met before cells lose their ability to self-renew and fully commit to differentiation. The outcome of Wnt9b reception is inherently dependent on the combined expression patterns of multiple proteins, including secreted agonists/co factors provided in trans, coupling all the NPC within the niche (Figure 5).

**Figure 5.**
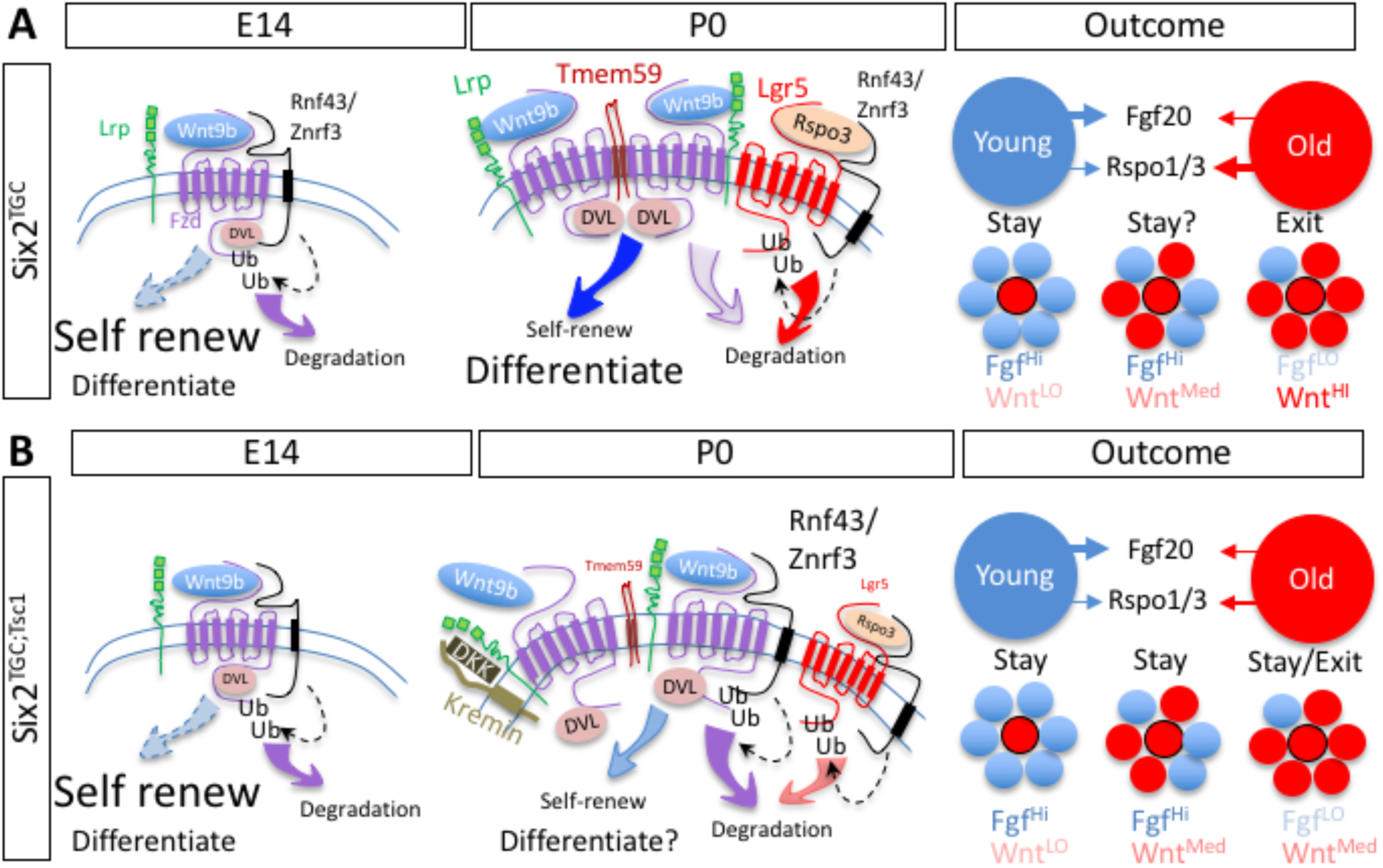
A parsimonious hypothesis of how NPC orchestrate the cessation of Nephrogenesis. (A) In E14 Six2^TGC^ controls, important Wnt agonist are transcribed but poorly translated, resulting in lower Wnt signal intensity and preference for self-renewal. Transcripts levels for the niche-retention signal Fgf20 are high (Chen et al., 2015). At P0, increased translation of Tmem59, Rspo1/3, and additional factors increase the stability of Fzd/Wnt complexes and promote signalosome formation. Wnt signal intensity now promotes differentiation over self-renewal. We assume other factors in the niche environment, including Fgf9/20, can modulate this in trans. If enough neighbors are in this state, an environment low in FGF and high in Rspo1/3 will drive the NPC to differentiate en masse. (B) In Six2^TGC; Tsc1^ cells, translation of Tmem59, Rspo1/3 is less efficient, and that of antagonists DKK, Kremen and RNF43 is more efficient. As a result, cells in the population are experiencing lower Wnt signals, delaying coordinated secession by an unknown amount sufficient to produce more nephrons. Experimental data suggest this delay is ∼12 hours. This environment may also increased the number of nephrons forming during arcading (the period during which the entire population remaining in the niche differentiates, (Rumballe et al., 2011)).

The implication of this study to human health is clear: it demonstrates that a subtle reduction in Wnt signaling can increase nephron endowment, expanding the possible therapeutic approaches to address clinical need in treatment of premature infants in addition to minimizing nephrotoxic insults. Similar observation was reported with a BMP antagonist, Decorin (Fetting et al., 2014). Given the centrality of Wnt and BMP to ongoing neurological developmental processes, they might not be the ideal target pathways, though this remains to be explored. Importantly, *Six2* hemizygotes NPC are distinct from *Tsc1* hemizygote NPC, suggesting that additional pathways operating at different time points all contribute to regulating nephron endowment, only some through extension of cessation timing or enhancement of late stage nephrogenesis. Of note, a later-acting mechanism may reflect a more targetable period of nephrogenesis for therapy in premature infants as more organ system complete their development. Furthermore, future investigation into the impact of Hamartin on translation may identify safer targets.

## MATERIALS AND METHODS

### Mouse Strains and Embryo Staging

The following lines were used in this study: *Tg(Six2-EGFP/cre)1*^*Amc*^ (Jax Stock No: 009600; herein *Six2*^*TGC*^), *Six2-<u>C</u>re<u>E</u>RT2*, (herein Six2^KI^) (Kobayashi et al., 2008), *Tsc1*^*f/f*^ (Kwiatkowski et al., 2002), B6; 129S4 - Gt(ROSA)26Sortm9(EGFP/Rpl10a)Amc/J (Jax Stock No: 024750; herein EGFP/Rpl10a), Rspo3^f/f^ (Jax Stock No: 027313; Rspo3^tm1.1Jcob^), and Foxd1^tm1(GFP/cre)Amc^ (herein Foxd1^Cre^; Jax Stock No: 012463). In mouse experiments using embryos, embryonic day (E) 0.5 was designated as noon on the day a mating plug was observed. All mice were maintained in the Cincinnati Children’s Hospital Medical Center (CCHMC) animal facility according to the animal care regulations. Our experimental protocols (IACUC2016-0022/0032 and IACUC2018-0107/0108) are approved by the Animal Studies Committee of CCHMC. Animals were housed in a controlled environment with a 12-h light/12-h dark cycle, with free access to water and a standard chow diet.

### Generation of TMEM59 Knockout mice

The methods for the design of guide RNA and the production of animals were described previously (Yuan et al., 2017). We targeted Tmem59 with a guide RNA (target sequence: ACGACCTCACCAGAGTCAGA) according to the on- and off-target scores from the web tool CRISPOR (http://crispor.tefor.net) (Haeussler et al., 2016). To form chimeric guide RNA, 5 ul crRNA (500 ng/ul; Integrated DNA Technologies (IDT), Coralville, IA) and 10 ul tracrRNA (500 ng/ul; IDT) were mixed in the IDTE buffer and heated to 95C for 2 min, followed by slow cooling to 25C. To form ribonucleoprotein complexes (RNPs), 3 ul chimeric guide RNA and 3 ul Cas9 protein (1,000 ng/ul; IDT) were added to 9 ul Opti-MEM and incubated at 37C for 15 min. The zygotes from super-ovulated female mice on the CD-1 genetic background were electroporated with 7.5 ul RNPs on ice using a Genome Editor electroporator (BEX Co. Ltd, Tokyo, Japan); 30V, 1ms width, and 5 pulses with 1s interval. Two minutes after electroporation, zygotes were moved into 500 ul cold M2 medium (Sigma-Aldrich, St. Louis, MO), warmed up to room temperature, and then transferred into the oviductal ampulla of pseudo-pregnant CD-1 females. Pups were born and genotyped by PCR and Sanger sequencing.

### Single-cell sample preparation and sequencing

E14 and P0 kidneys were isolated in ice-cold PBS. The capsule was removed, four (P0) or six (E14) kidneys were placed in 1.5ml Eppendorf, digested using 250ul of 10mg/ml *Bacillus licheniformis* cold-protease (Sigma-Aldrich) and incubated at 10°C in an Eppendorf Thermomixer R shaking at 1400rpm. After 5-7 minutes, the kidneys were removed, and the single-cell suspension was rinsed twice with 750ul of ice-cold 0.04%BSA/PBS. Trypan Blue stained cells were counted using a hemocytometer, requiring greater than 85% cell viability. Cells were resuspended at a concentration of 1000 cells/ul and loaded on a chromium 10x Single Cell Chip (10x Genomics, Pleasanton, CA). Libraries were prepared using Chromium Single Cell Library kit V2 (10x Genomics) and sequenced on an Illumina HiSeq-2500 using 75bp paired-end sequencing.

### Nephron progenitor cell isolation for translating ribosome affinity purification (TRAP)

To perform ribosome affinity purification on nephron progenitor cells (NPCs), timed-matings between *Six2*^*TGC*^ or *Six2*^*TGC;Tsc1*^ and EGFP-L10a mice were set, and embryos harvested at E14 or P0. Kidneys were isolated in ice-cold PBS and the capsule removed. Approximately 4-6 kidneys were placed in 1.5ml Eppendorf and digested with 250ul of 0.25%Trypsin/0.2% Collagenase (Sigma-Aldrich) at 20°C in a Thermomixer R. After 10-15 minutes, the kidneys were removed, and the single-cell suspension was rinsed twice with 750ul of ice-cold 0.3%BSA/PBS. NPCs were labeled using cell-surface primary antibodies (1/250 biotin-conjugated anti-ITGA8, R&D Systems, Minneapolis, MN; 1/250 PE-conjugated anti-PDGFRA, Invitrogen, Carlsbad, CA) while rocking on ice for 30-60 minutes. Cells were rinsed twice in 750ul of ice-cold 0.3%BSA/PBS and subsequently stained with Streptavidin APC (Invitrogen) while rocking on ice for 30-60 minutes. After two washes in 750ul of ice-cold 0.3%BSA/PBS, the NPCs were isolated by FACS on a SH800 cell sorter (Sony Biotechnology Inc., San Jose, CA) selecting for GFP^HI^ (Six2 and EGFP/Rpl10a-expressing NPC), ITGA8+ (NPC with some Stroma labeling, but not nephron epithelia, which remain EGFP/Rpl10a+) and PDGFRA- (Stroma is PDGFRA+) cells. Cells were collected in ice-cold 0.3%BSA/PBS. All procedures were performed with solutions containing 100ug/ml cycloheximide. Purification of polysome-bound RNA from kidneys was performed as previously described (Heiman et al., 2008). Briefly, 250,000-300,000 NPCs were homogenized in ice-cold lysis buffer (20 mM HEPES, 5 mM MgCl2, 150 mM KCl, 0.5 mM DTT, 1% NP-40, 100mg/ml cycloheximide, RNase inhibitors and protease inhibitors) by vigorous pipetting. The samples were centrifuged at 2,000 x g for 10 min at 4°C to remove large debris. Supernatants were extracted with 1% NP-40 and 30mM 1,2-diheptanoyl-sn-glycero-3-phosphocholine (DHPC, Avanti Polar Lipids, Alabaster, AL) on ice for 5min, and centrifuged at 20,000 x g. The clarified supernatant was incubated in low salt-buffer (20mM HEPES, 150mM KCl, 5mM MgCl2, 0.5mM DTT, 1% NP-40, RNAse inhibitors and 100ug/ml cycloheximide) containing streptavidin/protein L-coated Dynabeads (ThermoFisher, Waltham, MA) bound with 17μg of anti-GFP antibodies (HtzGFP-19F7 and HtzGFP-19C8, Memorial Sloan Kettering Centre, New York, NY) overnight at 4°C with gentle agitation. After 24 hours, the beads were collected using magnets for 1 minute on ice, and the beads washed four times in high-salt washing buffer (20mM HEPES, 350mM KCl, 5mM MgCl2, 0.5mM DTT, 1% NP-40, RNAse inhibitors and 100ug/ml cycloheximide) and collected with magnets. RNA was eluted from the samples and purified using a Qiagen (Hilden, Germany) microRNeasy RNA kit per manufacturer’s instructions. RNA integrity was assessed using an RNA PicoChip (Agilent Bioanalyzer, Santa Clara, CA) and the Clontech SMARTer library kit V2 (Takara Bio USA, Mountain View, CA) was used for cDNA library construction and sequenced on an Illumina HiSeq-2500 using 75bp paired-end sequencing.

### Total Protein quantification by Flow cytometry

Briefly, kidneys from E14 or P0, P1, or P2 mice were isolated in ice-cold PBS and the capsule removed. Four to six kidneys were digested with 1mg/ml neutral-protease and incubated at 20°C for ∼5-10 minutes in Thermomixer at 1400RPM. Since Six2TGC mice were used in these experiments, the kidneys were examined under a fluorescent microscope to ensure removal of nephron progenitors (GFP+ cells). It is important to note that isolating NPCs for flow cytometry using other enzymatic means, such as cold-protease trypsin, trypsin or collagenase had variable results. After digestion, the cells were washed twice with ice-cold 1%BSA/PBS, fixed with 4%PFA for 10 minutes and washed twice with ice-cold 1%BSA/0.1% Triton-100/PBS. Permeabilized cells were incubated with the primary antibodies (1/500 anti-GFP, Aves Labs, Tigard, OR; 1/250 anti-Tmem59, ProteinTech, Rosemont, IL; 1/250 anti-ITGA8, R&D Systems; 1/250 anti-PDGFRA-PE, Invitrogen) on ice for 60 minutes. After incubation, the cells were washed twice with ice-cold 1%BSA/0.1% Triton-100/PBS and stained with secondary antibodies (1/500 anti-Rabbit BV405, Jackson Immuno Research, West Grove, PA; 1/500 anti-Biotin APC, Jackson Immuno Research; and 1/500, anti-Chicken FITC, Jackson Immuno Research) for 30-60 minutes while rocking on ice. The cells were washed twice with ice-cold 1%BSA/0.1% Triton-100/PBS and the cells were analyzed on a BD FACS Canto II (BD Biosciences, San Jose, CA).

### Nephron counts

HCl maceration of whole kidneys was performed according to (MacKay et al., 1987; Peterson et al., 2019). Kidneys were isolated from ≥ P28 animals, the capsule removed, minced using a razor blade and incubated in the presence of 6N HCL for 90 min. The dissociated kidneys were vigorously pipetted every 30 minutes to further disrupt the kidneys. After incubation, 5 volumes of distilled water were added to the samples followed by incubation at 4°C overnight. 100ul of the macerate was then pipetted into a cell culture dish and glomeruli were counted in triplicate (three aliquots) for each sample. A single experimenter, blinded to the genotypes of kidneys being scored, performed all counts. Genotyping was performed after the counts on DNA isolated from the liver, spleen or tail using the primer sets reported as follows, 5’ – 3’: *Cre* (F-GCA TTACCGGTCGATGCAACGAGTGATGAG; R-GAGTGAACGAACCTGGTCGAAATCAGT GCG), *Tsc1* (F- AGGAGGCCTCTTCTGCTACC; R- CAGCTCCGACCATGAAGTG), and *Rspo3* (F-TATACTGCGATCTAATGCCCTCT; R-CCTTTCGACTTGATGGTGG).

### Translatome data analysis

RNA-seq data were analyzed using AltAnalyze using Kallisto to map transcripts to Ensmart72/mm10. Gene expression was computed as log2-transformed transcripts per million (TPM) and the significance of differential expression was set to a fold change of >1.25 and a raw p-value of <0.05. Visualization of the data was performed with Morpheus https://software.broadinstitute.org/morpheus/.

### Processing raw scRNA-Seq data

Raw scRNA-Seq data was sequenced using 10x Genomics v2 chemistry and was processed using Cell Ranger (v2.0.0) (https://github.com/10XGenomics/cellranger). Raw reads per cell were mapped to the mm10 mouse genome. As a quality control metric, barcoded cells with less than ∼5k UMIs were dropped from the analysis, and for 5 out of 6 samples, ∼66% of the reads were confidently mapped to the transcriptome with median of 2500 genes expressed per cell in each sample. Finally, mapped reads were used to generate matrix of raw counts of genes across barcodes for each sample. Please refer to the table for number of barcodes and percentage reads mapped per sample.

### Quality Filtering, dimension reduction and clustering

Seurat (v2.3.4) (https://doi.org/10.1016/j.cell.2019.05.031) package in R was deployed to perform quality control, selection of highly variable genes (HVGs), dimensionality reduction and clustering. Briefly, cells with at least 100 detected genes and any gene expressed in >= 3 cells were used in subsequent analyses. For quality control, cells with ∼3 fold higher than the median number of transcripts/cell (likely doublets) or cells with ∼3.5 fold higher than the median mitochondrial gene fraction were excluded. Log2 transform followed by scaling was performed to normalize the raw counts of genes across all cells. Using normalized counts, cell cycle effect on clustering was minimized by regressing the difference between G2M and S phase. Next, HVGs were selected by marking the outliers from dispersion vs. avgEXP plot. These HVGs were then used to perform principal component (PCA) analysis, a linear dimensionality reduction approach. Genes included in the first 20 principal components (PCs), were then used to cluster the cells, applying a graph-based clustering algorithm embedded in Seurat. Cell clusters were visualized using t-distributed stochastic neighbor embedding (tSNE) (a non-linear dimensionality reduction approach) and markers of each cluster were obtained using Wilcoxon rank-sum test in ‘FindAllMarkers’ function.

### Cell-type Annotation

Clusters were annotated to their respective cell-types using a priori knowledge of cell-type specific markers and a computational framework. Gene set enrichment analysis(GSEA; https://doi.org/10.1073/pnas.0506580102, https://www.nature.com/articles/ng1180) of our cluster markers against gene-sets defined in Combes et. al (Combes et al., 2019) assisted in the computational and statistical annotation of cell clusters. A matrix of false discovery rate (FDR) of enrichment of gene-sets in cell-clusters was extracted from GSEA results using a custom python script (available on GitHub).

### Integrating samples to understand cell-type specific responses

To determine cell-type similarities and differences among the samples, we implemented the integration analysis provided in Seurat (v3.0.0). Quality filtering, normalization and cell-cycle effect regression was carried out as described above. Instead of selecting outliers from dispersion vs avgExp plot, HVGs were determined using the variance stabilizing transformation “vst” function which yielded approximately 2000 HVGs. Integration of the samples was performed by first identifying “anchors” using ‘FindIntegrationAnchors’, followed by integrating the datasets using anchors with ‘IntegrateData’ function. Dimensionality reduction was performed by PCA followed by both uniform manifold approximation (UMAP) and t-SNE. Graph-based clustering was used to generate clusters. Seurat built-in functions were used to generate dotplots, feature plots and heatmaps. Photoshop were used to merge feature plots into the images shown in the figures.

### RNA-Velocity analysis

To analyze cell-state expression dynamics, we utilized the ‘run10x’ function in the velocyto program to compute the spliced to un-spliced ratios for each transcript in single cells (https://www.nature.com/articles/s41586-018-0414-6). For analytical purposes, a repeat masked gene transfer format file (gtf) was downloaded from UCSC and used in conjunction with a standard gtf from the Cell Ranger pipeline. The resulting file in ‘loom’ format was analyzed subsequently by scVelo - single-cell RNA velocity generalized to transient cell states (https://github.com/theislab/scvelo). Quality filtering, normalization, scaling and cell-cycle removal was carried out using the default parameters in scanpy package (https://github.com/theislab/scanpy). Velocity estimation was done using the ‘stochastic’ model to capture the steady states. The stochastic model is generated by treating transcription, splicing and degradation as probabilistic events, which aids in approximating steady state levels not only from mRNA levels but from inherent expression variability among the cells.

### Functional and Pathway enrichment

To determine the biomarkers, pathways and transcription factors (TFs) enriched in our cell-types, we utilized GO-Elite (Zambon et al., 2012), which can identify a minimal non-redundant set of biological functions and pathways enriched for a particular set of genes or metabolites. Markers identified by our Seurat analysis for each cluster were also analyzed by GO-Elite using the following parameters: 1.96 as z-score cutoff for pruning ontology terms, Fisher Exact Test for over-representation analysis (ORA), 0.05 as permuted p-value, at least 3 genes changed and 1000 permutations for ORA.

### Data and. Software Availability

All computer codes and scripts [R scripts, custom Python/Perl scripts and software packages] have been uploaded to GitHub [Link]. All codes have been licensed with a general public license (GPL v3.0). The data analyzed in this publication have been deposited in NCBI’s Gene Expression Omnibus (Edgar et al., 2002) and are accessible through GEO Series accession number XXXX and will contain all processed bam files, raw counts of genes across barcodes/cells and cell annotations in the form of a metafile. All the parameters used in the analysis of the samples are reported in the scripts are available on GitHub

## ACKNOWLEGMENTS

We would like to thank Dr. Yueh-Chiang Hu and the Transgenic Animal and Genome Editing Core facility and the Research Flow Cytometry Core (supported by NIH S10OD023410) at Cincinnati Children’s Hospital Medical Center. We thank Dr. John Cobb for the generation of the Rspo3^*f/f*^ mouse line. We also acknowledge funding support from the National Institute of Diabetes and Digestive and Kidney Diseases (NIH DK106225 to R.K.) as well as generous support from an internal General Funds award at Cincinnati Children’s Hospital Medical Center.

## SUPPLEMENTARY FIGURES

**Supplementary Figure S1:**
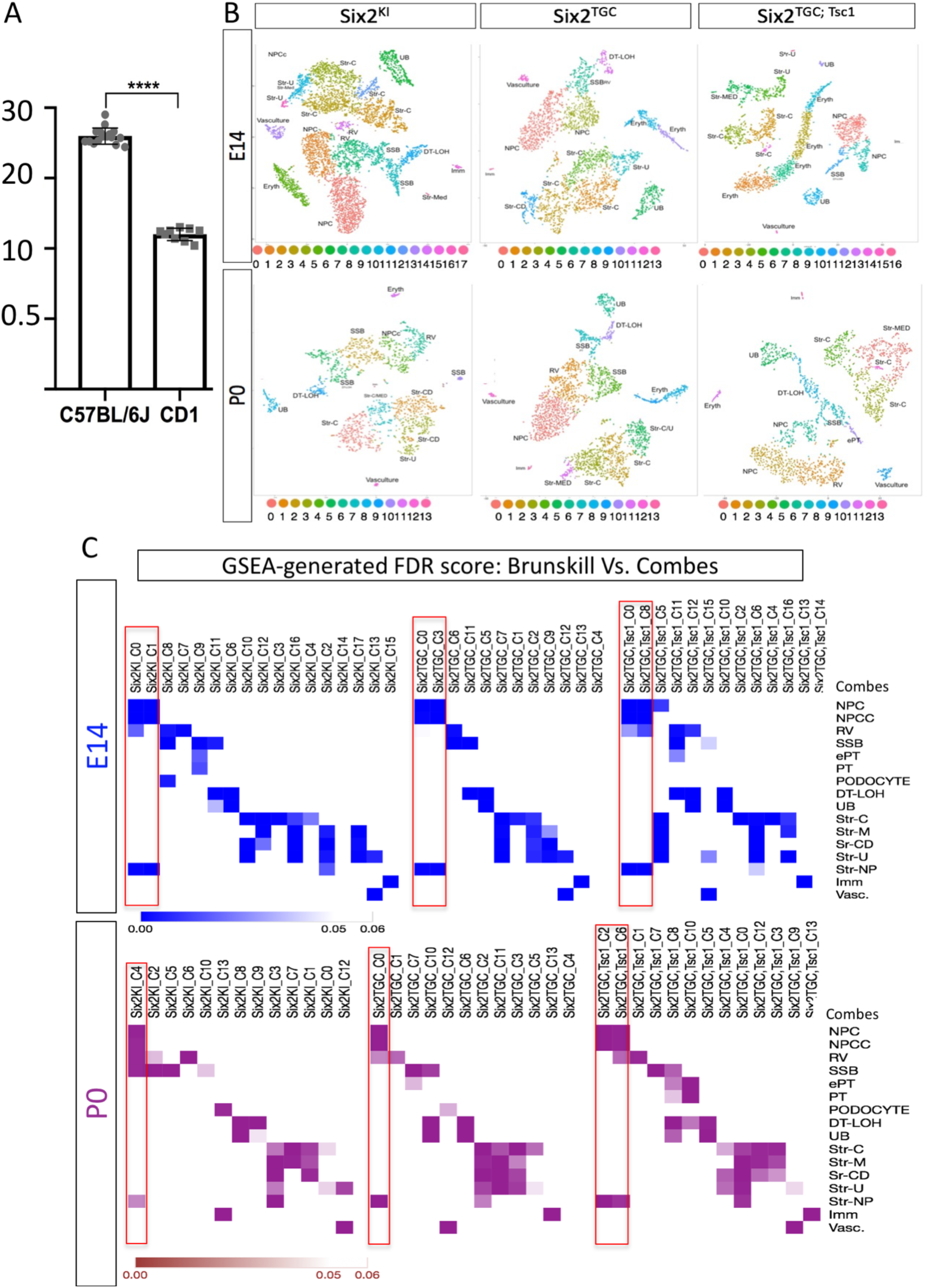
Nephron progenitor scRNA analysis. (A) Nephron endowment in the inbred strain C57BL/6J is nearly double that of the outbred strain CD1. Note the low intra-strain variance. Y-axis counts nephrons by the thousand (e.g., 10 is 10×10^3^). (B) tSNE analysis of six cortical samples from three strains and two developmental timepoints. The Six2^KI^ at E14 is analyzed in Figure 1. The clusters are colored, the key shown below each cluster. Each cluster was also named based on GSEA analysis. (C) GSEA analysis compared each of the clusters identified in (A) with the clusters identified by Combes et al. Best FDR value for each two-way comparison was extracted with a custom script (available in GitHub) to generate the similarity matrix shown. Note that only Cluster C5 in one of six samples (E14 Six2^TGC; Tsc1^) contained both stromal and NP markers and was excluded from downstream analyses. Clusters in the red boxes were used for the analysis shown in the text.

**Supplementary Figure S2:**
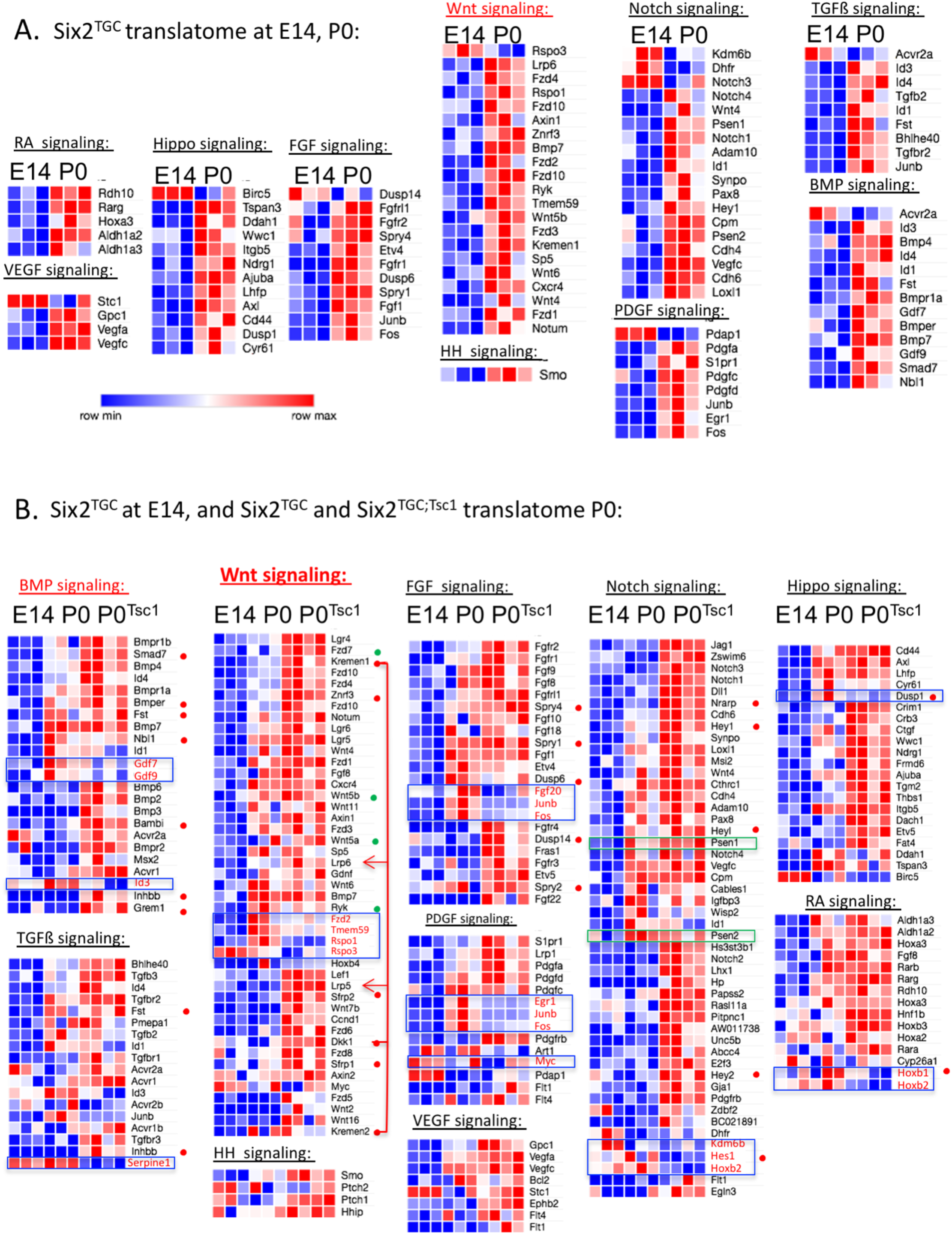
Analysis of signaling pathway components by TRAP. RNAseq analysis of polysomal transcripts is shown in supplementary Table S5. We selected transcripts based on membership in ten signaling pathways. The data were filtered to only show avg expression of Log2≥1; fold change >1.2 or <-1.2, and raw pValue <0.05 in at least one of the datasets in the comparison. (A) Transcripts that vary between E14 Six2^TGC^ and P0 Six2^TGC^, visualized to show high/low values within each row. (B) Transcripts that vary between E14 Six2^TGC^, P0 Six2^TGC^ and P0 Six2^TGC; Tsc1^, visualized to show high/low values within each row. Antagonists are marked by a red dot, non-canonical components marked by green dot. Transcripts differentially translated in *Tsc1* hemizygotes are enclosed in a blue box and marked with red text.DKK1 and Kremen protein target Lrp4/5/6 for degradation, shown with red lines and arrows. Note that the catalytic subunit in g-secretase, required to produce Notch signals (Psen1, Psen2), are expressed poorly in E14 (green boxes).

**Supplementary Figure S3:**
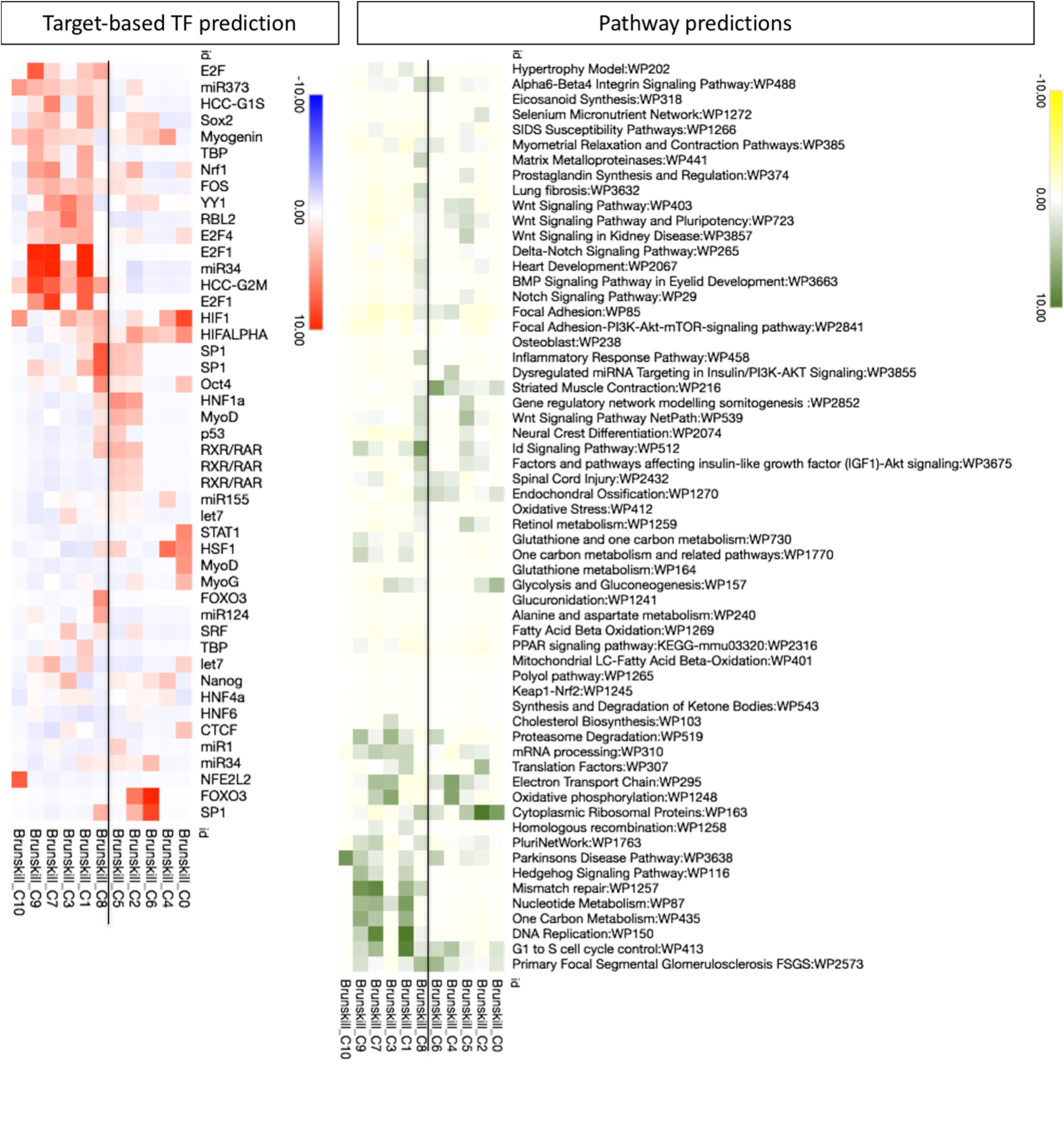
GoElite analysis. Transcripts from the boxed clusters in Supplementary Figure 1C were analyzed in GoElite to identify transcription factors likely to regulate the differential transcripts defining each cluster (Target-Based TF prediction) and signaling pathways likely to be active (Pathway Predictions). The clusters on the left of the vertical line are committed to differentiation, on the right, identified as cap mesenchyme. See text for details.

